# Rhizosphere Microbiome Influence on Tomato Growth under Low-Nutrient Settings

**DOI:** 10.1101/2024.08.13.607683

**Authors:** Gerardo Mejia, Angélica Jara-Servin, Luis Romero-Chora, Cristóbal Hernández-Álvarez, Mariana Peimbert, Rocío Cruz-Ortega, Luis D. Alcaraz

## Abstract

Studies have shown that reduced nutrient availability enhances microbial diversity around plant roots, positively impacting plant productivity. However, the specific contributions of rhizosphere microbiomes in nutrient-poor environments still need to be better understood. This study investigates the role of Plant Growth-Promoting Rhizobacteria (PGPR) in enhancing the growth of *Solanum lycopersicum* under hydroponic conditions. We hypothesised that nutrient limitation would increase the selection of beneficial bacterial communities, compensating for nutrient deficiencies. Our hydroponic system, with treatments consisting of 50% reduced fertiliser application supplemented with a soil-derived inoculum, exhibited greater bacterial diversity and biomass than controls, suggesting a successful enrichment of PGPR that compensates for nutrient deficiencies. Using 16S rRNA gene sequencing, we found a significant enrichment (*p* ≤ 0.001) and correlation with beneficial plant traits (*p* ≤ 0.05) of bacterial genera such as *Luteolibacter*, *Sphingopyxis*, and *Kaistia*. Shotgun metagenomics identified the critical role of *Methyloversatilis* in nitrogen fixation and other key taxa bacterial proteins in plant-bacteria interactions. Additionally, our findings identify core taxa across different cultivation systems. These results support the potential for microbiome engineering to enhance microbial diversity and plant productivity, offering a path to reduce fertiliser use in agriculture and improve sustainability.

## Introduction

The tomato (*Solanum lycopersicum* L.), domesticated in Mexico and Peru, is a widely consumed staple with high nutritional value, and its cultivation area has doubled over the past two decades (Jaiswal, et al., 2020; Tieman, 2017). Beyond its agricultural importance, the tomato serves as a model organism in scientific research due to its sequenced genome, diverse genetic tools, short life cycle, and role in studying plant-microbe interactions (van Rengs et al., 2022; Adedayo et al., 2022). A key concept in plant-bacteria interactions is the two-step model for microbiome acquisition, which describes the mutual selection between plants and soil microorganisms, shaping the rhizosphere and endosphere microbiomes (Bulgarelli et al., 2013; Sasse et al., 2018). This model has been validated in tomatoes, revealing a core microbiome influenced by diverse soil geochemistries (Barajas et al., 2020).

The interaction between the tomato microbiome and host selection is reciprocal, as specific bacterial genera, such as *Streptomyces*, *Bacillus*, *Pseudomonas*, and *Flavobacterium*, associate with tomato quantitative trait loci (QTLs) and participate in essential metabolic functions like iron and sulfur metabolism and vitamin synthesis (Oyserman et al., 2022). Soilless and hydroponic cultivation are practical experimental approaches due to reduced pathogen risk (Anzalone et al., 2022). Microorganisms in hydroponic conditions primarily originate from plants, seeds, water, insects, and personnel; they generally show less variation than soil microbiomes (Escobar et al., 2021; Anzalone et al., 2022).

The host plant’s nutrition significantly influences its microbiome. Plants rely on soil microbes for essential nutrients in nutrient-poor conditions, increasing microbial diversity to aid nutrient acquisition. Microbes play crucial roles in nitrogen fixation, phosphorus mobilisation, and overall plant health, as shown in nutrient-stressed soybean and maise studies (van der Heijden et al., 2008; Meier et al., 2022; Wang et al., 2024). Agricultural practices, including chemical fertilisation, can deplete plant-available nutrients and reduce below-ground diversity, leading to land degradation and environmental pollution (Pahalvi et al., 2021). Over-application of nitrogen fertilisers can cause soil acidification, increasing toxic aluminium levels that inhibit root growth (Kopittke et al., 2019).

The success of PGPR inoculants for crops depends on factors like bacterial colonisation of plant roots, root exudation, and environmental conditions, all of which can affect microbial inoculation outcomes (Santoyo et al., 2021). While initial microbial community differences may arise from different fertilisers, plant development strongly influences microbial community structures over time (Grunert et al., 2019; Naumova et al., 2022). The relationship between microbial diversity and plant productivity, particularly in nutrient-limited conditions, requires further investigation.

We hypothesise that low-nutrient environments increase selection pressure for root-associated bacteria, potentially recruiting a PGPR community that compensates for nutrient scarcity. This study aims to identify and characterise these PGPR communities and their genetic traits in *S. lycopersicum* under nutrient-limited conditions, providing insights into their adaptation and contribution to plant health.

## Materials and methods

### Soil sampling used as inoculants

We collected seven distinct soil samples to explore the potential for recruiting PGPR communities. Sampling locations were selected based on vegetation types, including forests, grasslands, and agricultural areas. Each 2 kg sample was stored in sterile bags and refrigerated at 4°C until inoculation experiments began. Metadata for each site included the collection date, geographic coordinates, altitude, and observations documented (Table S1).

The soil samples were used as microbial inoculants in a hydroponic system with two controls: one with 100% fertilisation and another with 50% fertilisation, both without soil (Fig. 1). The experimental design included two treatments: non-sterilised soil was directly inoculated into the hydroponic system, and sterilised soil was used to assess nutritional contributions without native microbiota (Nutrient Control, NC).

**Figure 1.**
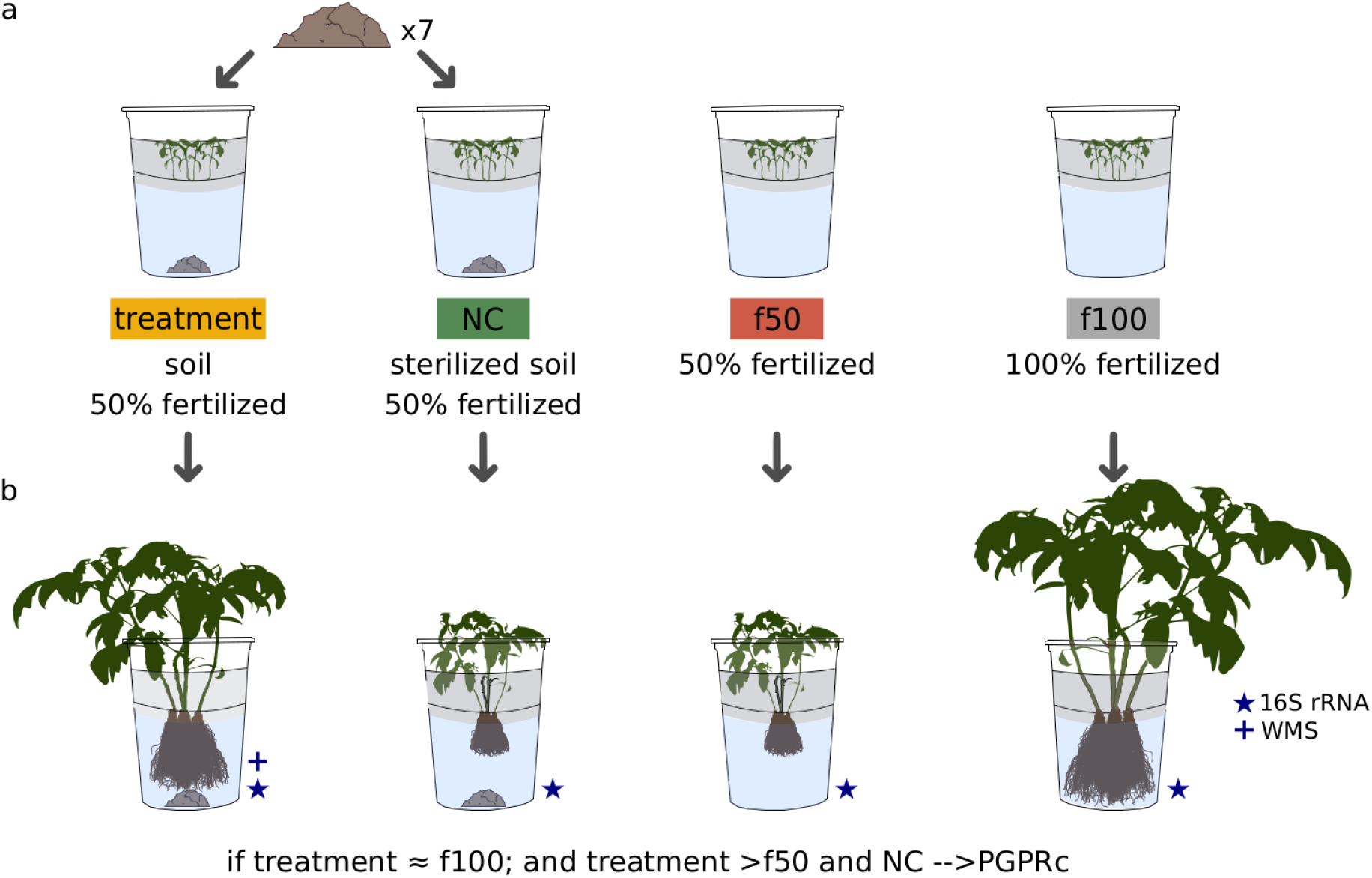
Experimental Layout. The diagram summarizes the methods to evaluate the impact of PGPR communities and fertilisation levels in tomato (*Solanum lycopersicum*) hydroponic system: a) Treatment Groups (yellow): Non-sterilized soil with microbial inoculants and 50% fertilisation; seven different soils were tested individually. Nutrient Control (NC, green): Sterilized soil with 50% fertilisation. 50% Fertiliser Control (f50, red): Reduced nutrient conditions without microbial inoculation. 100% Fertiliser Control (f100, grey): Full nutrient conditions without microbial inoculation. b) We selected inoculated plants with non-sterilized soil that showed better growth under 50% fertilisation than sterilised soil and f50, highlighting the beneficial effects of microbial communities on plant growth. WMS: Whole Metagenome Sequencing.

We hypothesised that if growth-promoting communities are present, plants inoculated with non-sterilised soil will grow better with 50% fertilisation than those with sterilised soil, indicating beneficial microbial communities (Fig. 1).

### Plant-growth conditions

We regulated nutrient concentration and mechanical soil properties using a hydroponic system with plastic cups and nylon netting containers (Alatorre-Cobos et al., 2014). The cups were sterilised with 70% ethanol and 4.5% commercial sodium hypochlorite. *Solanum lycopersicum* L. cv. Rio Grande seeds were surface-sterilised with 70% ethanol for one minute, followed by a 2.5% NaClO wash for two minutes, and rinsed with sterile distilled water. The seeds were germinated in chambers for 72 hours at 21°C with a 12-hour light/dark cycle before being transferred to hydroponic containers, with five plants per container. The plants were grown for 60 days in a greenhouse under natural light and dark conditions.

We compared plant phenotypic traits—stem length, dry weight, stem diameter, and leaf area—between treatments and controls to identify significant growth improvements. Each treatment included seven triplicate soil samples and three triplicate controls, with 5 g of soil added to a medium containing 50% fertiliser. Controls consisted of a non-inoculated set with 50% fertiliser (f50) and another with 100% fertiliser (f100) to evaluate growth under varied nutrient conditions. Nutrient controls (NC) were established by autoclaving the inoculum twice (15 minutes at 121°C, 15 psi, with a 24-hour interval), and these received 50% fertilisation (Fig. 1). The experiment used 240 plants across treatments and controls. We evaluated the potential of the seven collected soils to recruit PGPR, testing 15 plants per soil type. Based on plant phenotypes, we selected three soils with potential PGPR effects for further analysis using Whole Metagenome Shotgun and 16S amplicon sequencing. The experiment used 240 plants across treatments and controls. We evaluated the potential of the seven collected soils to recruit PGPR, testing 15 plants per soil type. Based on plant phenotypes, we selected three soils with potential PGPR effects for further analysis using Whole Metagenome Shotgun and 16S amplicon sequencing.

The hydroponic medium for the f100 treatment contained per litre: 0.5g MgSO _4_, 0.45g KNO_3_, 0.22g KH_2_PO_4_, 0.8g citric acid, 0.1g EDDHA-Fe, 1g Ca(NO_3_)_2_, and 0.3g STEM (S 13%, B 1.35%, Cu 2.3%, Fe 7.5%, Mn 8%, Mo 0.04%, Zn 4.5%). The treatments, f50, and NC used a medium with half of these nutrient concentrations (Table S2).

Phenotypic traits were measured at the end of the experiment. Stem length was measured from the base to the apical meristem (Ohta & Ikeda, 2015). Stem diameter was measured below the cotyledons with an electronic Vernier (Jwa & Walling, 2001). Leaf area was estimated using aerial photographs analysed with GIMP (GIMP development team, 2021) and ImageJ (Schneider et al., 2012). Plants were oven-dried at 45°C for ten days to determine dry weight.

### Metagenomic DNA Processing, Sequencing, and Analyses

Metagenomic DNA was extracted from rhizosphere and endosphere fractions using the Mobio PowerSoil DNA extraction kit (MoBio, Carlsbad, CA, USA) following Lundberg et al. (2012). For 16S rRNA gene sequencing, the V3-V4 regions were amplified (Herlemann et al., 2011) and sequenced on an Illumina MiSeq platform. Shotgun metagenomic sequencing of both fractions was conducted using the Illumina NextSeq 2x150 at the LANGEBIO Genomic Services Laboratory.

The 16S rRNA gene was amplified using 341F/805R primers targeting the V3-V4 regions, following the Illumina MiSeq protocol with 5’ overhangs. Duplicate PCR reactions for each sample were performed with Pfx platinum polymerase under the following conditions: 95°C for 3 min; 5 cycles at 94°C for 30 s, 55°C for 30 s, 68°C for 30 s; then 25 cycles at 94°C for 5 s and 68°C for 30 s; and a final extension at 68°C for 5 min. Products were purified with the SV Wizard PCR Purification kit (Promega, WI, USA).

Detailed bioinformatics and statistical analysis procedures for 16S sequencing are available on GitHub (genomica-fciencias-unam/tomato-hydroponics). Briefly, raw reads were trimmed using the FASTX-Toolkit (’fastx_trimmer’) (Hannon, 2010) when the Phred quality score was below 20. Forward reads were trimmed to 250 bases, and reverse reads to 220 bases. The reads were assembled with pandaseq (Masella et al., 2012) using a minimum overlap of 250 bp and a quality threshold 0.6. Operational taxonomic units (OTUs) were defined at 97% identity and clustered using cd-hit-est v4.7 (Li & Godzik, 2006). Singleton sequences, chimaeras (detected by blast_fragments method (Caporaso et al., 2010), mitochondria, and chloroplast sequences were removed. Taxonomy was assigned using BLAST (Altschul et al., 1990) with the SILVA v138.1 database (Quast et al., 2013).

Metagenomic reads were quality-filtered with Trimmomatic v0.36 (Bolger et al., 2014). Sequences matching the *S. lycopersicum* genome (NCBI BioProject: PRJNA66163) were removed with Bowtie2 v2.3.4.1 (Langmead & Salzberg, 2012). Quality-filtered sequences were assembled with metaSPADES v3.12.0 (Nurk et al., 2017). Unmapped reads were reassembled with Velvet v1.2.10 using a k-mer size 31 (Zerbino & Birney, 2008). Contigs from both assemblies were used to predict ORFs with Prodigal v2.6.3 (Hyatt et al., 2010). Predicted genes were annotated against the M5nr database (Wilke et al., 2012) using Diamond v0.9.18.119 (Buchfink et al., 2015). Taxonomic assignment of predicted proteins was performed with Kraken2 v2.1.2 (Wood et al., 2019) using default parameters and the PlusPF database with RefSeq indexes (O’Leary et al., 2016). An abundance table was generated by mapping reads against predicted proteins using Bowtie2. Unannotated proteins were clustered with cd-hit v4.7 (Li & Godzik, 2006) at 70% identity.

Rhizosphere metagenome data from soil-grown tomatoes of the same variety (*S. lycopersicum* var. Río Grande) were retrieved from NCBI BioProject: PRJNA603603 (Barajas et al., 2020) and processed alongside the current sequencing data. For comparative analysis, both datasets were compared using subsamples of each metagenome by selecting 1 x 10^6^ random reads in three replicates.

### Identification of the PGPR-Associated Taxa and Metagenomic Comparative Analysis

We analysed bacterial taxa in treated and control groups using the 16S rRNA gene. We calculated relative abundances and employed Pearson’s correlation to examine relationships with plant phenotypic variables, focusing on taxa consistently present with high correlation coefficients (r² > 0.7).

Using the 16S dataset, we distinguished hydroponic core microbiome by analysing genus intersections across treatment samples.

To understand soil’s role in shaping the root microbiome, soil-grown tomato microbiomes were characterised using the 16S rRNA gene, following Barajas et al. (2020).

Hydroponic inocula originated from agricultural soils, differing from ruderal soils used for soil-grown tomatoes, though both shared the same plant genotype. Samples were re-analysed to identify the core microbiome by determining all samples’ genera intersections. We defined a Hydroponic Tomato Core Metagenome (HTCMe) from protein families shared across all hydroponic treatments. At the same time, the Tomato Core Metagenome (TCMe) was established from protein families in all rhizosphere samples from both conditions (hydroponics and soil). Non-metric dimensional scaling (NMDS) was used to compare protein families across both datasets.

### Key Hydroponic Bacteria Pangenome Analysis

The 16S amplicon analysis identified genera with high correlation values to plant traits, overrepresented in treatments and present in the core microbiome. We searched for their pangenomes in the hydroponic rhizosphere metagenomes. Using Prokka v1.12, we annotated reference genomes from selected bacterial species: *Flavobacterium* (43 genomes), *Hyphomicrobium* (6), *Luteolibacter* (8), *Methyloversatilis* (3), and *Sphingopyxis* (19), chosen for their association with tomatoes and prevalence in our samples. Roary software calculated pangenomes aligned with hydroponic metagenomic reads using Promer v3.23. Alignment percentage identity and coverage were visualised with identity graphs created using the promer_deid_v9gmv.py script (available on GitHub).

### Data analyses

Statistical analyses of plant phenotypes were conducted using R v3.5.1 (R Development Core Team, 2003), with ggplot2 v3.3.0 (Wickham, 2016) for plot design. Diversity analysis utilised R’s phyloseq v1.24.2 (McMurdie & Holmes, 2013). Correlations between OTU abundance and plant phenotype were analysed using corrplot v0.84 (Wei, 2021). Genera analysis in hydroponics and core microbiome calculations employed the Venn diagram v1.6.20 (Chen, 2022). Protein core metagenome was determined with UpSet v1.4.0 (Conway, 2017). Differential taxa and predicted proteins were calculated using DESeq2 v1.10.1 (Love et al., 2014) in R v3.2.2. CAP analysis used the Bray–Curtis dissimilarity score, constrained by plant phenotypic traits. NMDS ordination used Bray–Curtis dissimilarity scores between predicted proteins of tomato hydroponic and soil systems.

## Results

### Plant productivity and microbiome diversity under low nutrient availability

We hypothesise that nutrient limitation enhances the selection of PGPR bacterial communities, aiding plants in compensating for nutrient deficiencies. To test this, we used *S. lycopersicum* plants in a hydroponic system. Treatments involved plants with a low nutrient profile (50% fertilisation) and soil as a microbial inoculum. Controls included a group without soil or inoculum but with 50% fertiliser (f50) and another with 100% fertiliser (f100) to evaluate growth under different nutrient conditions. Nutrient controls (NC) were prepared with sterilised inoculum and received 50% fertilisation (Fig. 1).

Our assessment of several plant phenotypes revealed the effects of PGPR communities. The highest average biomass was recorded in the maximum fertilisation control (f100 = 4.86 g ± 0.71), followed by the 50% fertilised pots with soil as microbial inoculants (treatment = 2.77 g ± 0.46). Sterilised soil with 50% nutrients produced lower biomass (NC = 1.87 g ± 0.57), while the minimum fertilisation control (f50) produced the least biomass (0.83 g ± 0.45). These findings underscore the potential of PGPR bacteria in enhancing plant growth, particularly under nutrient-limited conditions.

Biomass production in treatments significantly exceeded that of the f50 and NC groups, demonstrating enhanced performance (t-test, *p* ≤ 0.001 and *p* ≤ 0.05, respectively; Fig. 2a and b). Similarly, the highest stem diameters were observed in f100 (5.60 mm ± 0.29) and treatments (4.81 mm ± 0.16), with treatments showing significant improvements over NC (3.70 mm ± 0.39) and f50 (3.12 mm ± 0.72) (t-test, p ≤ 0.001 and p ≤ 0.5, respectively). The stem length of treatments (32.84 cm ± 2.34) was comparable to f100 (37.16 cm ± 4.64), showing no significant difference, despite being significantly longer than those of f50 (20.7 cm ± 4.15) and NC (26.99 cm ± 2.85) (t-test, p ≤ 0.05 and *p* ≤ 0.01, respectively).

**Figure 2.**
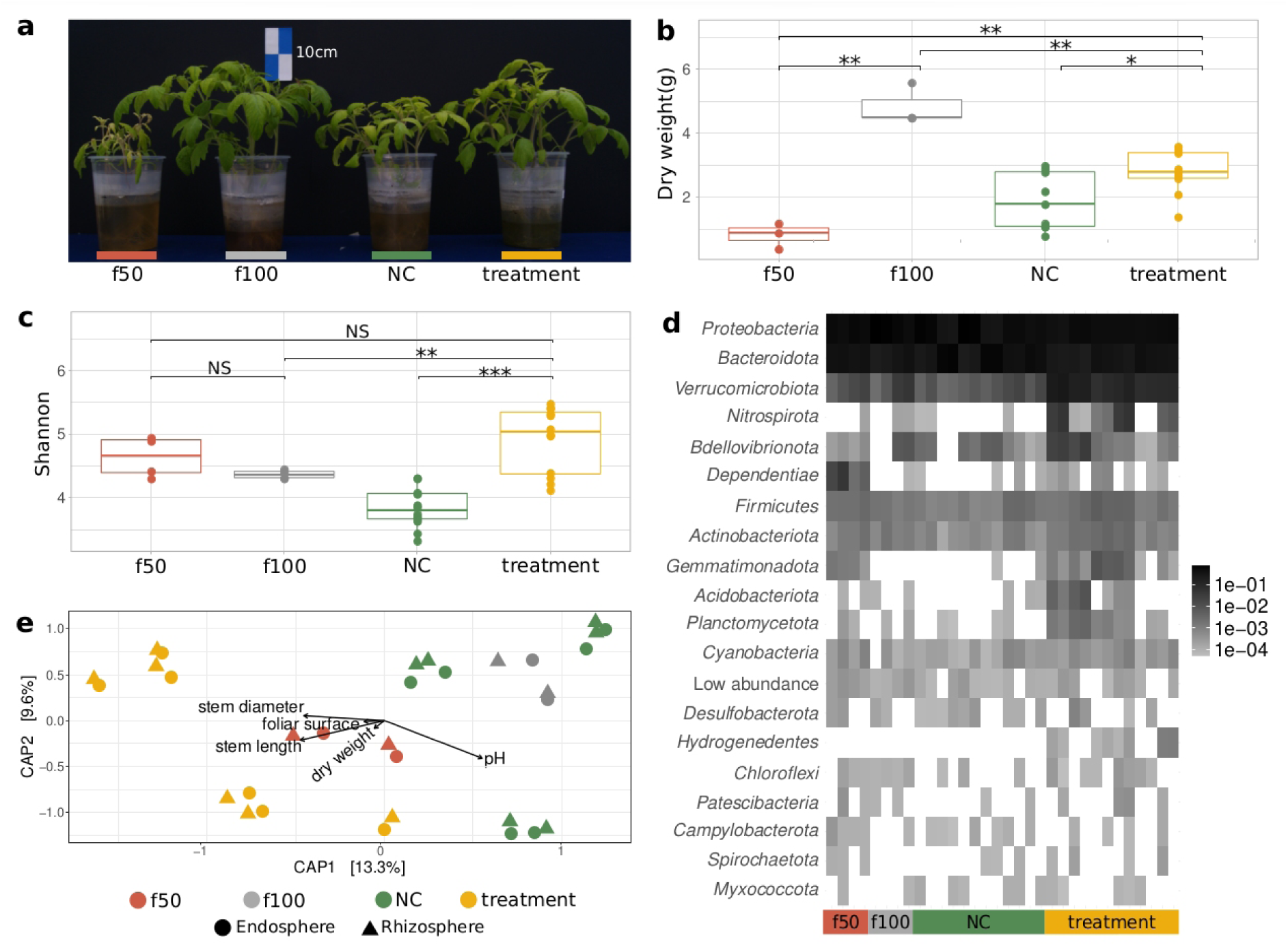
Impact of PGPR on Plant Diversity and Growth. (a) Representative plants from the PGPR community search system: 50% fertiliser control (red, f50), 100% fertiliser control (grey, f100), nutrient control (green, NC), and treatment (yellow), illustrating visual differences in plant growth. (b) Box plots showing the Shannon diversity index across treatment groups, indicating variations in microbial diversity. (c) Box plots of plant biomass production, highlighting growth differences under varying nutrient and microbial conditions (*\*p* < 0.05, \**\*p* < 0.01, \*\*\**p* < 0.001). (d) Heatmap depicting the relative abundance of phyla within each treatment and control group, with phyla of low relative abundance (≤ 0.001) shown in the “Low abundance” category. (e) CAP ordination plots based on Bray-Curtis distances, correlating operational taxonomic units (OTUs) with plant phenotypic traits and pH across all groups, illustrating the influence of microbial diversity on plant characteristics.

No significant differences were observed in leaf area between treatments (1390.79 cm² ± 211.21) and NC (1072.62 cm² ± 336.20) or between treatments and f50 (690.93 cm² ± 451.71). Additionally, wet weights were similar across treatments (10.59 g ± 1.47), f50 (5.86 g ± 3.41), and NC (9.24 g ± 2.18).

The substantial growth observed in treatments, compared to f50 and NC, suggests a beneficial impact of PGPR communities (Fig. 2a and Table S3). In contrast, chlorosis’s low performance and symptoms in the f50 group indicate nutritional stress and suboptimal growth conditions (Fig. 2a).

### Plant root diversity under low nutrient availability

We sequenced 3,591,762 reads (average = 284 bp) across 32 samples. After quality control and sequence clustering, we identified 12,294 Operational Taxonomic Units (OTUs; 97% 16S rRNA gene identity). These OTUs were classified within the hydroponic dataset, encompassing treatments, nutrient controls (NC), and fertilised controls (f100 and f50).

Treatments exhibited the highest richness with 7,247 OTUs (Chao1 = 3,223.19 ± 321.24), followed by NC with 5,265 OTUs (Chao1 = 2,812.76 ± 268.88), f50 with 3,077 OTUs (Chao1 = 3019.89 ± 666.25), and f100 with 2,957 OTUs (Chao1 = 2812.95 ± 323.14). The Shannon index revealed that treatments had the highest bacterial diversity (median = 5.03 ± 0.51), followed by f50 (4.66 ± 0.32). f100 and NC showed reduced diversity (4.36 ± 0.07; 3.80 ± 0.31, respectively). Significant differences were found in the diversity of treatments compared to all other groups (ANOVA, *p* = 3.48 x 10^-6^; Tukey-HSD, *p-adj* < 0.05), indicating a robust microbial enrichment under treatment conditions (Fig. 2b).

### Bacteria found in tomato roots under low nutrient availability

Taxonomic assignments of OTUs revealed that Proteobacteria was the dominant Phylum across all conditions. However, treatments displayed a lower proportion of Proteobacteria (average = 49.66%) compared to f100 (66.71%), f50 (55.35%), and NC (52.05%). Bacteroidota followed in abundance, with treatments showing a lower percentage (33.09%) compared to f50 (40.89%) and NC (46.54%), while f100 had the lowest (30.16%). Verrucomicrobiota was notably more abundant in treatments (13.59%) than all controls, indicating distinct microbial community structuring under treatment conditions (Fig. 2d).

The 7,247 OTUs were assigned to 818 bacterial genera. *Flavobacterium* was the most abundant in treatments (x̄ = 22.79 %) and had a substantial relative abundance growth in NC (x̄ = 45.14%). In contrast, the relative abundance of *Flavobacterium* was lower in f100 (x̄ = 18.65 %) and f50 (x̄ = 16.33 %). On the other hand, *Allorhizobium*-*Neorhizobium*-*Pararhizobium*-*Rhizobium (ANPR),* considered a single genus by the SILVA database, was the most prevalent genus in f100 (x̄ = 29.70%) and f50 (x̄ = 19.13%). However, rhizobia relative abundance decreased in both treatments (x̄ = 6.49%) and NC (x̄ = 14.66%). Notably, *Luteolibacter* showed a remarkable increase in treatments (x̄ = 11.76%) when compared with NC (x̄ = 0.09%), f100 (x̄ = 0.02%), and f50 (x̄ = 0.009%) (Fig. S1, Table S4, and S5).

### Potential PGPR associated with tomato under low-nutrient conditions

A Constrained Analysis of Principal Coordinates (CAP) was performed on the 16S communities using Bray-Curtis distances (Fig. 2e), explaining 22.9% of the variance in microbial communities, with CAP1 accounting for 13.3% and CAP2 for 9.6%. The treatment groups formed distinct clusters, separate from the controls. Significant differences were found between the treatments, NC, and controls (*p* = 1 x 10^−4^; Adonis), with controls (f50 and f100) showing greater dispersion. A notable difference was observed between nutrient controls (NC; sterilised soils with 50% fertiliser) and treatments (soil inoculum with 50% fertiliser), indicating distinct microbial community differentiation. Plant phenotypic traits such as stem diameter, leaf area, and medium pH significantly influenced microbial community structure, with treatment communities being more similar than any control.

Phenotypic traits and microbial genera abundance helped identify bacteria associated with PGPR activity. Significant Spearman’s correlations (*p* ≤ 0.05) showed that genera strongly correlated with beneficial plant phenotypes were prevalent in treatments. In contrast, those with low or negative correlations were found in NC and f50 controls (Fig. 3a). Genera such as *Sphingopyxis, Hyphomicrobium, Bradyrhizobium, Neorhizobium, Flavihumibacter, Luteolibacter*, and *Sphingobium* showed the highest correlation with positive plant traits (Table S6). Qualitative and quantitative analyses delineated shared and unique bacterial genera between treatments and controls. A Venn diagram revealed 179 unique genera within treatment groups, suggesting a rich diversity of potentially beneficial microbes (Fig. 3b, Table S7). We found unique genera in treatments: *Kaistia*, *Peredibacter*, *Nitrosococcus*, *Leucobacter*, *Pseudoruegeria*, and *Desulfonispora*. Additionally, these genus were positively correlated with plant phenotypes (Fig. 3a).

**Figure 3.**
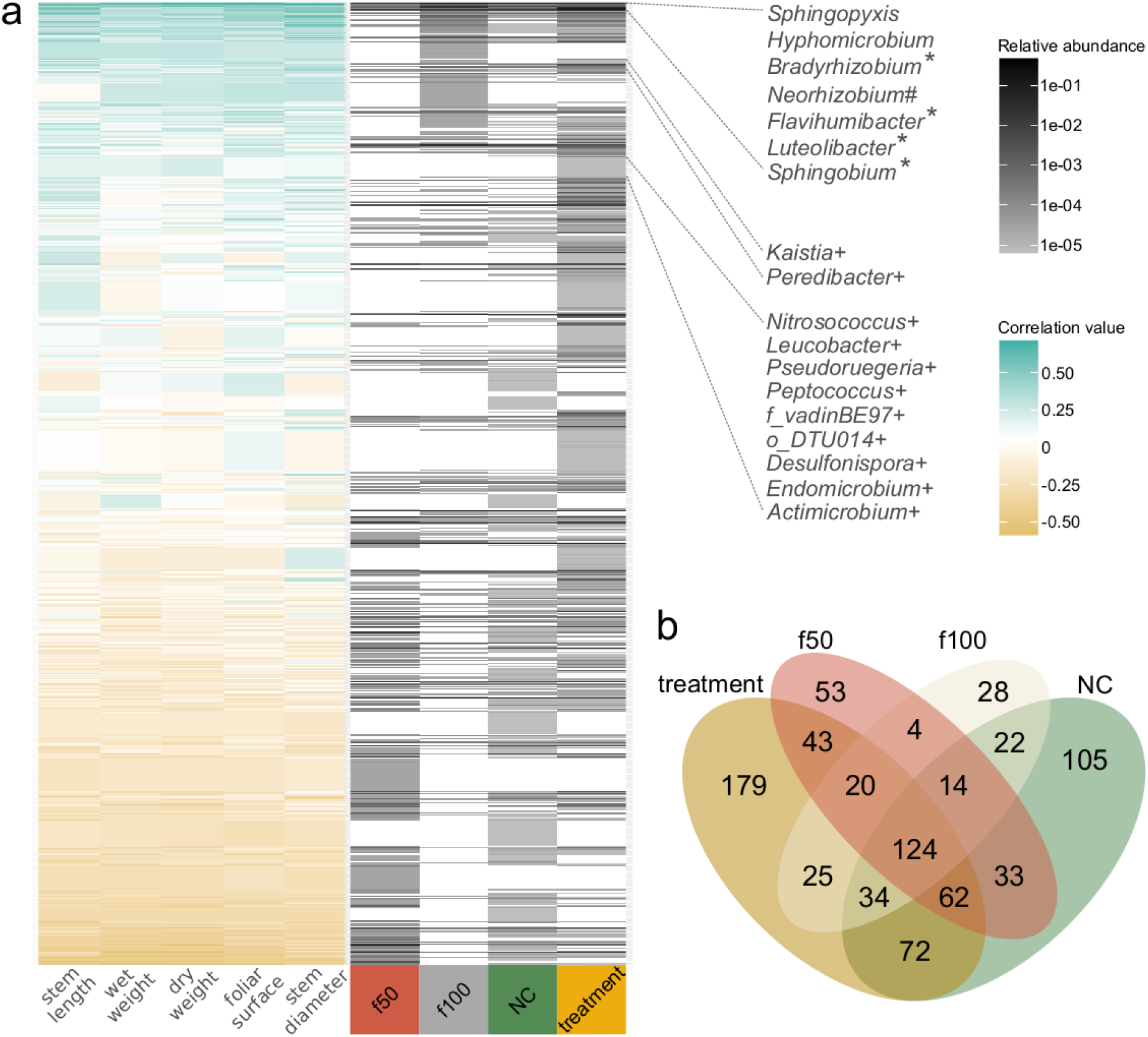
Relationship Between Microbial Genera and Plant Phenotypes. (a) Heatmaps showing the correlation between microbial genera relative abundance (grey scale) and plant phenotypic traits across treatment and control groups. Genera are sorted by decreasing correlation values, indicating their association with plant growth characteristics. Symbols denote genera overrepresented in treatments (*), controls (#), and those unique to treatments (+). (b) Venn diagram illustrating the distribution of microbial genera in controls and treatments, highlighting shared and unique communities to emphasise differences in microbial recruitment and establishment under varying conditions.

Sixty-six genera showed significant abundance variations in treatments (p-adj *≤ 0.05, DESeq2), including Caedibacter, Luteolibacter, Flavihumibacter, Sphingobium*, and *Bradyrhizobium*, associated with positive plant traits (Table S8). Additionally, 13 genera showed significant changes in controls, such as Brevundimonas, the combined group *Allorhizobium-Neorhizobium-Pararhizobium-Rhizobium*, *Phenylobacterium*, and *Flavobacterium*, also positively correlated with plant phenotypes (Table S6). This analysis highlights the dynamic interactions between microbial communities and plant phenotypic traits under varying nutrient conditions, emphasising the potential of PGPR in enhancing plant growth and resilience.

### Shared core taxa of tomato microbiome under hydroponics and soil growth

Our research identified a core microbiome in hydroponic tomato rhizospheres, consistent across all treatments and comprising 52 genera, collectively known as the Hydroponic Tomato Core Microbiome (HTCM; Fig. 4a). Key genera include *Flavobacterium*, *Luteolibacter*, *Sphingobium*, *ANPR*, *Caedibacter*, *Rhodobacter*, *Dyadobacter*, and *Sphingopyxis*. We also identified a relaxed core featuring genera present in at least 11 of the 12 treatment samples, such as *Flavihumibacter*, *Porphyrobacter*, *Defluviicoccus*, *Edaphobaculum*, *Hyphomonas*, *Gemmatimonas*, *Candidatus Protochlamydia*, *Bradyrhizobium, and Nordella* (Fig. 4a).

**Figure 4.**
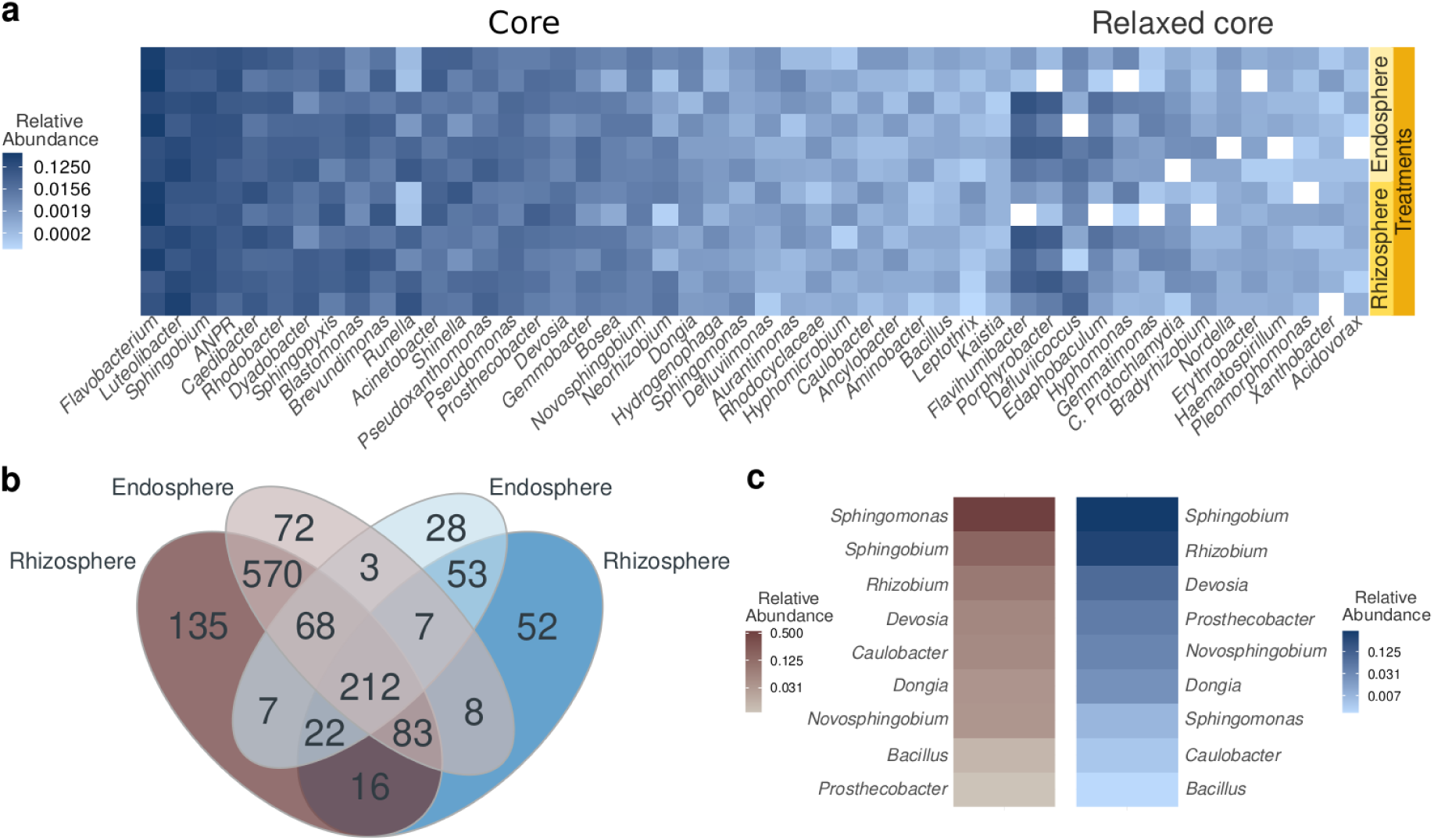
Analysis of the Tomato Core Microbiome in Hydroponic and Comparison With Soil Systems. a) Heatmap showing the relative abundances of genera within the Hydroponic Tomato Core Microbiome (HTCM). Relaxed core was determined by accounting for the presence of each genus in all samples except one. ANPR (*Allorhizobium Neorhizobium Pararhizobium Rhizobium*). b) The Venn diagram shows the shared genera between tomatoes in soil and hydroponics. c) Relative abundance of genera found in a strict core for hydroponic and soil rhizosphere samples.

Comparing hydroponic and soil-grown tomato roots revealed 212 common genera, 133 unique to hydroponics, and 777 exclusive to soil (Fig. 4b, Table S9). The HTCM and Soil Tomato Core Microbiome (STCM) comparison showed 87 genera in soil rhizospheres and endospheres, with nine genera common to both systems, forming the Hydroponic-Soil Tomato Core Microbiome (HSTCM). This hybrid core includes *Sphingomonas*, *Sphingobium*, *Rhizobium*, *Devosia*, *Caulobacter*, *Dongia*, *Novosphingobium*, *Bacillus*, and *Prosthecobacter* (Fig. 4c).

The relative abundance of these genera varied, with *Sphingobium* more prevalent in hydroponics and *Sphingomonas* more abundant in soil, highlighting their adaptability and crucial role in the tomato root microbiome across different conditions.

### Tomato root metagenomic diversity under low nutrient availability

Hydroponic treatment metagenomes generated 126,030,926 raw reads. After quality control and filtering out host plant sequences (*Solanum lycopersicum* L.), 114,475,444 sequences remained, assembling into 5,239,742 contigs with an average length (N50) of 1,506 base pairs. Protein prediction identified 5,196,182 proteins grouped into 1,984,648 protein families at 70% identity. Of these, 722,677 proteins were annotated using the M5nr database (Table S10).

Notable proteins in hydroponic treatments included the IS110-like element ISPa71 transposase (0.05% relative abundance) and others such as hypothetical proteins hp_5547 (0.037%) and hp_304 (0.028%), a DUF11 domain-containing protein (0.023%), glycosyltransferase (0.022%), and a Z1 domain-containing protein (0.020%). Proteins involved in heavy metal efflux, like the CusA/CzcA family RND transporter, also showed a 0.020% abundance (Table S11).

Non-metric dimensional scaling (NMDS) analysis showed distinct clustering of predicted proteins of hydroponic and soil-grown tomato metagenomic proteins (Fig. S2a). Hydroponic samples contained significantly more proteins (334,724) than soil samples (142,737; *p* ≤ 0.01 Wilcoxon). The Shannon-Weaver diversity index was higher in hydroponic samples (H’ = 11.62 ± 0.22) compared to soil samples (H’ = 10.97 ± 0.28) (Table S12, Fig. S3), likely due to higher sequencing coverage in hydroponics (42 x 10^6^ reads vs. 29 x 10^6^ in soil).

DESeq2 analysis identified significant differences between protein families in hydroponic (1,388 families) and soil metagenomes (330 families) (*p-adj* ≤ 0.001) (Fig. S2b). Proteins like the IS110-like element ISPa71 transposase and the TonB- dependent receptor were more prevalent in hydroponic samples (Table S13). Cluster of Orthologous Groups (COG) categorisation showed abundant categories in hydroponics: replication, recombination, and repair (L COG); unknown functions (S COG); amino acid transport and metabolism (E COG); translation, ribosomal structure, and biogenesis (J COG); and inorganic ion transport and metabolism (P COG) (Fig. S2c).

*Luteolibacter* was the most prevalent genus in hydroponic taxonomy, with significant representation from *Luteolibacter luteus* (12.14%) and *Luteolibacter ambystomatis* (11.9%). Other notable classifications included Bacteria (4.37%), Proteobacteria (1.61%), *Pseudomonas mexicana* (1.53%), and *Methyloversatilis* sp. RAC08 (0.93%) (Table S14, Fig. S4).

### Tomato Core Metagenome Analysis

We identified a core set of 48,116 protein families in all hydroponic tomato treatments, termed the Hydroponic Tomato Core Metagenome (HTCMe) (Fig. S5a, Table S15). The HTCMe includes various proteins such as the IS110-like element ISPa71 family transposase, DUF11 domain-containing protein, glycosyltransferase, Z1 domain-containing protein, CusA/CzcA family heavy metal efflux RND transporter, and PAS domain S-box protein. Notably, it also contains proteins involved in plant growth promotion, including tRNA dimethylallyl transferase MiaA, indole acetamide hydrolase, tryptophan decarboxylase, aldehyde dehydrogenase, and components of the nitrogenase enzyme complex (*nifD, nifK, nifH, nifA, nifB, nifE, nifN, nifW,* and *nifZ*).

Using metagenomic data from soil-grown tomatoes (Barajas et al., 2020), we established the Soil-grown Tomato Core Metagenome (STCMe) (Fig. S5b, Table S16). We also defined the Tomato Core Metagenome (TCMe) as protein families common to all soil-grown and hydroponic tomato rhizosphere samples. The TCMe comprises 663 protein families, including BamA/TamA family outer membrane proteins, TamB domain-containing proteins, patatin-like proteins, aspartate aminotransferase family proteins, TonB-dependent receptors, NAD-glutamate dehydrogenase, and glutamate synthase large subunit. These proteins were consistently detected across all samples, underscoring their fundamental role in tomato physiology across different growing conditions (Fig. S5c, Table S17).

### Key Bacteria Identification in Metagenomes

We constructed genus-level pangenomes using the reference genomes of *Luteolibacter*, *Flavobacterium*, *Sphingopyxis*, and *Hyphomicrobium*, which were identified as key taxa through 16S rRNA analysis. These pangenomes served as anchors to recruit metagenomic reads, verifying their presence in the hydroponic rhizosphere. Comparison with hydroponic metagenomic reads showed high similarity, with amino acid identities ranging from 70% to 100%, confirming their involvement in the hydroponic rhizosphere (Fig. 5a).

**Figure 5.**
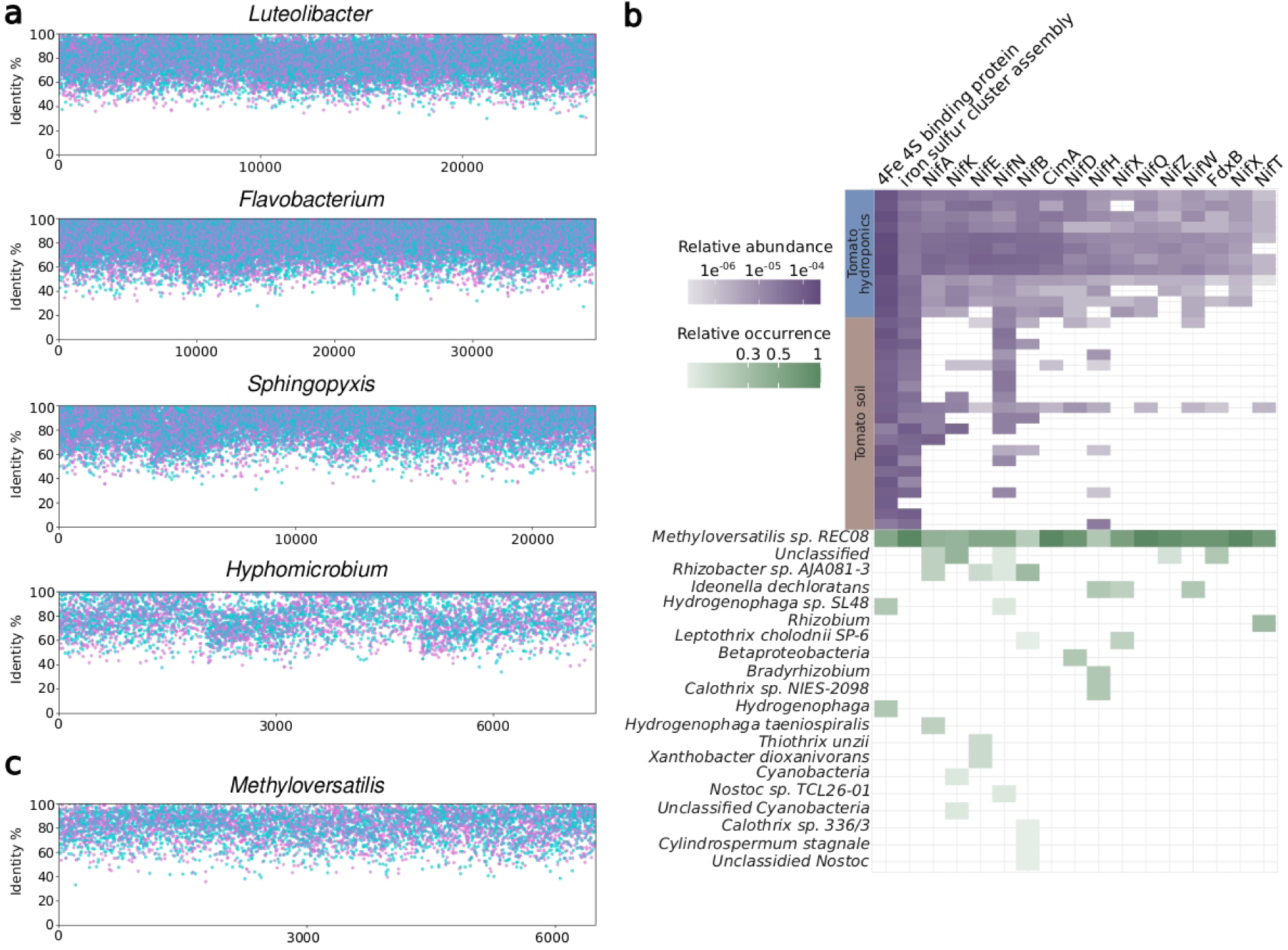
Pangenomic Analysis of Key Microbial Genera in Tomato Rhizospheres. a) Recruitment plots showing hydroponic rhizosphere metagenomes six-frame translations and its amino acid identity against *Luteolibacter, Flavobacterium, Sphingopyxis*, and *Hyphomicrobium* pangenomes. Blue dots are forward and pink ones reverse matches. b) Heatmap of nitrogen fixation proteins showcasing the relative abundance in hydroponic (blue axis) versus soil (brown axis) tomato metagenomes. A green heatmap is also included to present a taxonomic revision of each protein, detailing the taxa associated with each nitrogen fixation protein and their relative abundances. c) Coverage and average amino acid identity graphs for *Methyloversatilis*. This panel highlights the genomic representation and sequence conservation of *Methyloversatilis* in the metagenomes, underscoring its significance in nitrogen metabolism within the hydroponic system.

We also identified nitrogen fixation-related protein families using Level 4 SEED categories, showing an increased abundance in hydroponic treatments compared to soil metagenomes, especially the nitrogenase complex (Fig. 5b). Taxonomic identification with Kraken2 revealed that the *Methyloversatilis* genome contained all identified Nif-related proteins, followed by unclassified Cyanobacteria and *Rhizobacter* sp. AJA081-3. An analysis of the *Methyloversatilis* pangenome confirmed its presence in hydroponic metagenomes, with amino acid identity percentages ranging from 70% to 100% (Fig. 5c).

## Discussion

### Microbiome diversity and plant growth

As expected, the highest microbiome diversity was observed in treatments (Fig. 2c), while control groups f50, f100, and the nutrient control NC group showed lower diversity due to the lack of additional microbiological input. However, the Shannon diversity values in our treatments (4.92 ± 0.29) were lower than those reported in previous studies of soil-grown tomatoes. Cai et al. (2017) reported an average diversity of 5.38, and Barajas et al. (2020) reported values between 6.21 and 7.75. In hydroponic conditions, Shannon values ranged from 3.2 to 4.3, as Anzalone et al. (2022) noted, suggesting that the confined environment of hydroponics limits microorganism movement toward plant roots, reducing diversity. Anzalone et al. (2022) also reported a decrease in fungi in hydroponics compared to soil-grown tomatoes.

The fertilised controls showed predictable growth patterns, with the f100 controls achieving the highest biomass, indicating their nutritional needs were fully met. However, they exhibited lower bacterial diversity than the f50 controls (Fig. 2c), supporting van der Heijden et al. (2008), who hypothesised that optimal nutrient conditions reduce the need for plants to recruit a diverse microbial community. Treatments with 50% fertilisation and inoculation showed greater bacterial diversity and biomass than the f50 and NC controls. This suggests that plants recruit PGPR under nutrient-restricted conditions to support growth. Our findings indicate that reduced nutrient availability increases the recruitment of beneficial microbial diversity, enhancing plant productivity. Similar results have been reported in rice and soybean under low-nutrient conditions (Sinong et al., 2021; Wang et al., 2024) and in maise, where the root microbiome is linked to plant genetic variability under low nitrogen conditions (Meier et al., 2022).

The variation in bacterial communities between treatments and controls (Fig. 2e) demonstrates a link between the microbiome and plant phenotype, indicating that our system facilitates the recruitment of rhizosphere communities from the inoculum and tracks those promoting plant growth. Our study found a high proportion of the phyla Proteobacteria and Bacteroidota in both control and treatment groups (Fig. 2d, Table S4), consistent with previous reports on tomatoes (Cai et al., 2017; Zolti et al., 2019). These phyla and Actinobacteriota are typically dominant in the rhizospheres of tomatoes and other *Solanum* spp. (Zolti et al., 2018; Barajas, 2020).

Verrucomicrobiota was the third most abundant phylum in the tomato rhizosphere, consistent with findings from hydroponic tomato systems (Vargas et al., 2021). In rice studies, Verrucomicrobiota has been linked to root growth-promoting traits (Bünger et al., 2020). Our research showed enrichment of Verrucomicrobiota in treated groups compared to controls, suggesting that this phylum may significantly promote plant growth under our hydroponic conditions.

*Luteolibacter*, a Verrucomicrobiota genus enriched in treatment groups, correlated positively with plant phenotypes and was part of the Hydroponic Core Microbiome (HTCM). Found in diverse environments like potato and leek rhizospheres and marine settings (da Rocha et al., 2010; Park et al., 2013), *Luteolibacter* has potential as a PGPR in heavy metal-contaminated environments (Zadel et al., 2020). Proteins related to stress responses, such as those protecting against reactive oxygen species, were identified in *Luteolibacter* (Table S5, S7, S13).

While *Flavobacterium* (Bacteroidota) was the most abundant genus across treatments and controls, it did not directly correlate with plant phenotypic traits (Tables S5 and S6), indicating a key microbiome role despite not directly affecting plant phenotype, aligning with previous observations (Anzalone et al., 2022).

*Sphingopyxis* (Alphaproteobacteria) demonstrated the highest positive correlations with plant traits (Fig. 3a) and was overrepresented in treatments (Table S8). Known for producing indoleacetic acid (IAA) and enhancing plant growth, its association with quantitative trait loci (QTLs) in tomatoes underscores its PGPR role (Oyserman et al., 2022).

*Hyphomicrobium* (Alphaproteobacteria), although not predominant, showed significant correlations with plant traits (Table S6). Its role in the tomato rhizosphere is likely enhanced by its facultative methylotrophy, allowing it to thrive in hydroponic systems by utilising methane (Martineau et al., 2013).

Genera exclusive to treatments, such as *Akkermansia*, *Methylocapsa*, *Arenibacter*, *Marimicrobium*, *Syntrophomonas*, *Nitrosococcus*, *Thiohalobacter*, and *Kaistia*, could be critical for plant growth. *Kaistia*, which correlated positively with plant traits and was identified in the HTCM, is known for promoting plant growth under phosphorus limitation (Liu et al., 2023). Metabolites from *Kaistia* have been reported to modulate the biofilm and motility of *Methylobacterium* (Usui et al., 2020a,b), which was also found in the HTCM and positively correlated with plant phenotype. These findings suggest that *Kaistia* and *Methylobacterium* might be relevant in promoting tomato growth.

### Proteins in hydroponic tomatoes under a low nutrient concentration

We identified 1,984,648 protein families with 70% identity in hydroponic treatments targeting plant growth promotion. Proteins overrepresented in hydroponic metagenomes spanned various COG categories, with ’Replication and Repair of DNA (L)’ being most prevalent (Fig. S2c). PGPR community bacteria likely use these proteins to repair DNA damage from reactive oxygen species (ROS) under low oxygen hydroponic conditions. Notable proteins included catalase, cytochrome C peroxidase, and Vanadium-dependent haloperoxidase, known for ROS protection. Chaperones ClpB and GroEL highlighted adaptation mechanisms crucial in oxygen-limited conditions and key to survival in hydroponics.

The Hydroponic Tomato Core Metagenome (HTCMe) comprised 48,116 protein families in all conditions, while the Tomato Core Metagenome (TCMe) included only 663 families. The most prevalent HTCMe protein, the IS110-like element ISPa71 family transposase, supports the hypothesis that the rhizosphere facilitates horizontal gene transfer (Maheshwari et al., 2017), allowing DNA rearrangements that help bacteria adapt to environmental changes (Lugtenberg & Dekkers, 1999). The HTCMe core features proteins directly related to plant growth, such as enzymes involved in nitrogen fixation (Table S15).

HTCMe includes the BamA/TamA family outer membrane protein and TamB domain. TamA and TamB are part of the translocation and assembly module (TAM) subunits, which assemble outer membrane proteins related to adhesion and biofilm formation in bacteria (Josts et al., 2017). These proteins are crucial for infection and host colonisation (Heinz et al., 2015), indicating TAM’s potential role in colonising tomato roots and other plants.

Our study also explored the pangenomes of microbial genera in the hydroponic root metagenomes, confirming their presence and identifying genes associated with plant growth promotion. For example, genes within *Luteolibacter* and *Sphingopyxis* contigs included those related to multidrug resistance efflux pumps and chaperones, which may protect bacteria from plant-derived antibacterial compounds (Paço et al., 2016; Alvarez-Ortega et al., 2013). This finding suggests that these efflux pumps play a crucial role in the successful colonisation and persistence of these bacteria in the hydroponic rhizosphere.

### Nitrogen fixation and assimilation in the recruited PGPR community

Studies have shown that nitrogen fixation genes are more abundant in our tomato hydroponic system than in grown-soil tomatoes, emphasising the role of microbial communities in enhancing plant growth under controlled conditions (Bulgarelli et al., 2013). For example, the nitrogenase-stabilizing protein NifW is crucial for maintaining nitrogenase activity in aerobic conditions, highlighting its significance for diazotrophic bacteria in oxygen-rich environments (Nonaka et al., 2019). The Mo-dependent nitrogenase, which converts nitrogen into ammonia, is enriched in these systems (Seefeldt et al., 2018). Also hydroponic enriched were the nitrogenase molybdenum iron protein subunits and NifB, NifE, and NifN, involved in the biosynthesis of the nitrogenase cofactor (Rettberg et al., 2020). These findings suggest that nitrogen fixation is a crucial mechanism by which the hydroponic microbiome supports plant growth, highlighting the microbial communities’ adaptability to hydroponic cultivation challenges.

Nitrite can be reduced to ammonia by ferredoxin-nitrite reductase (NirA) in *Luteolibacter* or by nitrite reductase NADPH large subunit (NasD) in *Luteolibacter, Sphingopyxis*, and *Methyloversatilis*. This involvement in the nitrogen cycle highlights the collaborative nature of microbial communities in facilitating plant nitrogen acquisition, which is essential for optimal growth. *Methyloversatilis* plays a unique role in the hydroponic system by potentially using methanol or methane for carbon and energy while contributing to nitrogen fixation (Doronina et al., 2014; Smalley et al., 2015). Methane in anoxic zones of the hydroponic rhizosphere may enable *Methyloversatilis* to perform functions such as nitrate reduction, illustrating the complex interplay of microbial activities that support plant growth (Sun et al., 2016).

This research highlights the importance of understanding microbial dynamics in hydroponic systems as a pathway to enhance agricultural productivity and sustainability. Excessive fertilisation can lead to nutrient pollution, greenhouse gas emissions, biodiversity loss, and health impacts. Our model shows reduced nutrient availability increases microbial diversity, enhancing plant productivity under controlled conditions. By leveraging microbiome engineering, we can reduce fertiliser use, recruit a robust PGPR community, and improve plant growth and resilience, making agricultural practices more sustainable and environmentally friendly.

## Acknowledgements

We thank Laura Patricia Olguín Santos for her technical support in the Tempered Greenhouse during our experimental phase. We also thank Viviana Escobar for her technical assistance in the laboratory. Hugo Barajas for his help with soil samplings. This work was supported by Universidad Nacional Autónoma de México by the project DGAPA-PAPIIT-UNAM IN206824 to LDA and a Conahcyt Ph.D. scholarship (CVU 817269) to GM.

## Author contributions

Gerardo Mejia: conceptualisation, data curation, experiments, formal analysis, investigation, methodology, visualisation, and writing–original draft. Angelica Jara- Servin: Data curation, methodology, and visualisation. Luis Romero-Chora: Methodology. Cristóbal Hernández-Álvarez: conceptualisation, and methodology. Mariana Peimbert: resources. Rocío Cruz-Ortega: conceptualisation, resources. Luis D. Alcaraz: conceptualisation, data curation, formal analysis, funding acquisition, investigation, methodology, project administration, resources, visualisation, and writing–original draft.

## Data availability

Datasets generated for this study can be found in NCBI Bioproject PRJNA984704. Amplicon 16S rRNA sequencing data is available from SRR24973180 to SRR24973211 accessions. Raw shotgun sequences are available from SRR24973791 to SRR24973793 accessions. The bioinformatic protocols are available on GitHub: <u>genomica-fciencias-unam/tomato-hydroponics</u>. Raw Data necessary for the analyses at FigShare: 10.6084/m9.figshare.26543761.

## Supporting Information

**Fig. S1. Genera relative abundance in treatments and controls**. Heatmap depicting the relative abundance of genera within each treatment and control group. (Low relative abundance ≤ 0.001).

**Fig. S2. Comparative Analysis of Metagenomic Predicted Proteins in Hydroponically and Soil-Grown.** a) A non-metric multidimensional scaling (NMDS) plot was constructed using Bray-Curtis distances, comparing protein family composition (both annotated and hypothetical) between hydroponic and soil tomato rhizospheres. This visualisation highlights the distinct protein profiles in each cultivation method. b) Volcano plot illustrating significant differences in protein families (log2FoldChange) between hydroponic and soil rhizosphere metagenomes. This plot identifies statistically overrepresented or underrepresented proteins, indicating potential functional adaptations to hydroponic conditions. c) Classification of Clusters of Orthologous Groups (COG) for proteins overrepresented in hydroponic tomato treatments compared to soil.

**Fig. S3. Metagenomic diversity in hydroponics and soil.** Box plots showing observed protein families and the Shannon and Simpson diversity indexes across hydroponic and soil-grown tomato metagenomes.

**Fig. S4. Protein classification taxonomy in hydroponic metagenomes.** Percentage of proteins assigned to each taxon (Low percentage ≤ 0.1).

**Fig. S5. Tomato core metagenome.** a) Upset plot illustrating the proteins shared by all hydroponic metagenomic samples. b) An upset plot shows the proteins found in all soil metagenome samples. c) Venn diagram comparing the hydroponic tomato core metagenome (HTCMe) and the soil-grown tomato core metagenome (STCMe).

Table S1. Soil collection metadata.

Table S2. Fertiliser content.

Table S3. Plant phenotypic variables.

Table S4. Genera abundance per sample.

Table S5. Genera relative abundance average.

Table S6. Correlation of genera relative abundance and phenotypic traits.

Table S7. Genera Venn list controls and treatment.

Table S8. DESeq2 differential genera of treatment against controls.

Table S9. Venn list of genera for hydroponic and soil tomatoes.

Table S10. Metagenome, reads, contigs, and protein families number.

Table S11. Abundance of predicted proteins in metagenomes.

Table S12. Diversity index of metagenome hydro and soil samples.

Table S13. DESeq2 differential proteins of hydroponic metagenome against the soil.

Table S14. Taxonomic classification of predicted proteins.

Table S15. Hydroponic Tomato Core Metagenome.

Table S16. Soil-grown Tomato Core Metagenome.

Table S17. Tomato core Metagenome.

