## Supplemental Fig. 1 for "Rhizosphere Microbiome Influence on Tomato Growth under Low-Nutrient Settings"

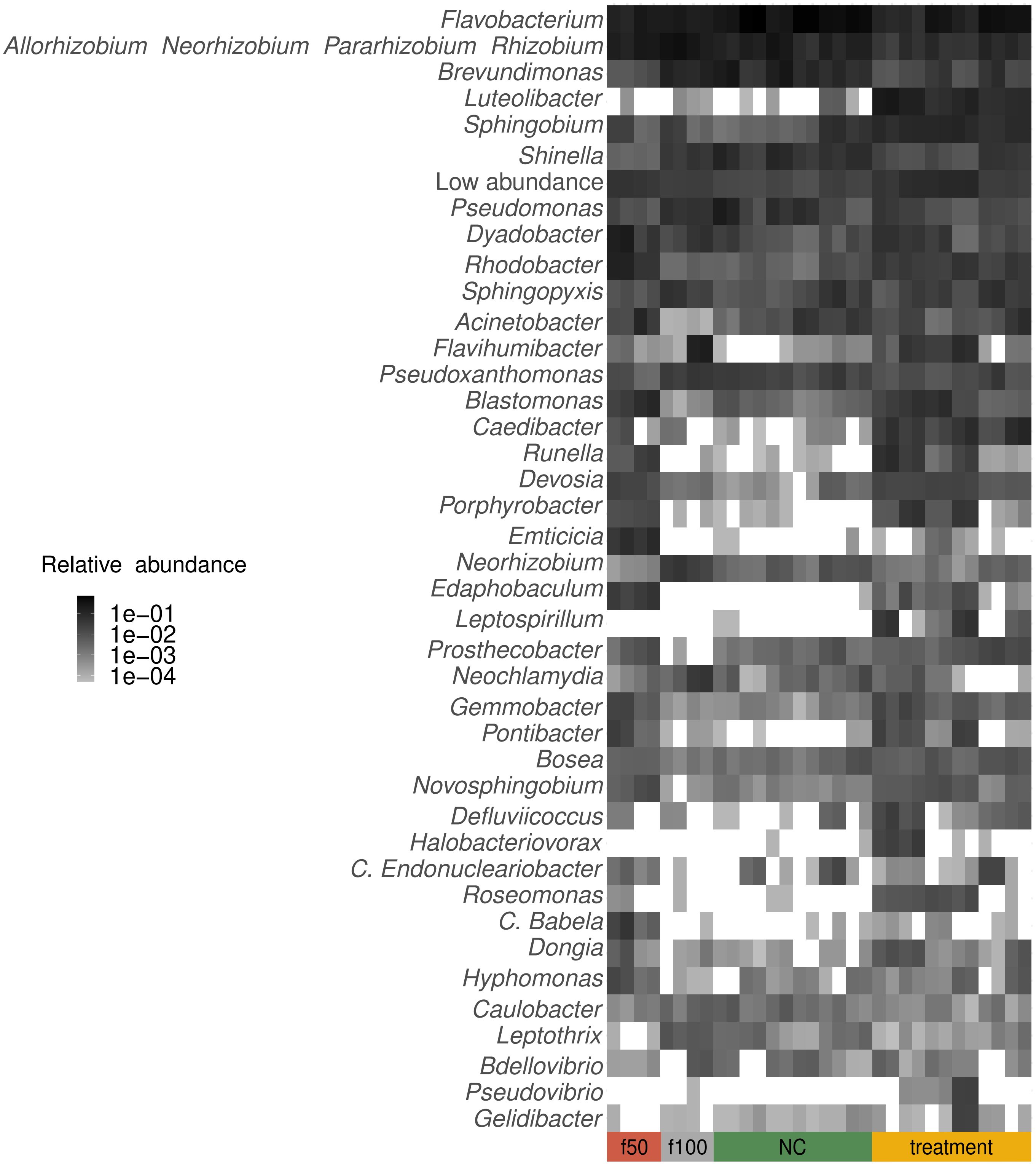


**Figure S1. Genera relative abundance in treatments and controls.** Heatmap depicting the relative abundance of genera within each treatment and control group. Low abundance relative abundance ≤ 0.001.
