## Supplemental Fig. 2 for "Rhizosphere Microbiome Influence on Tomato Growth under Low-Nutrient Settings"

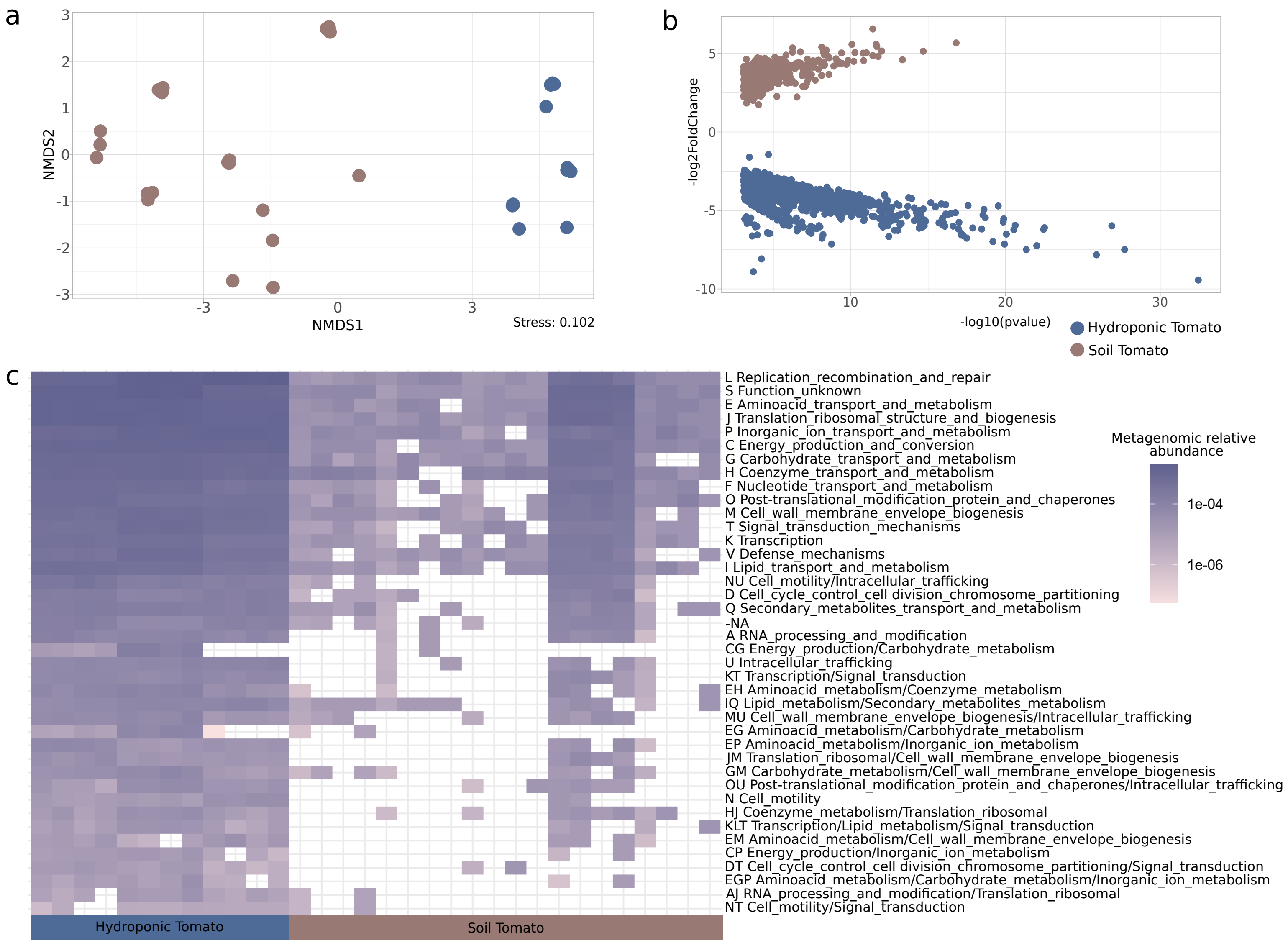


**Figure S2. Comparative Analysis of Metagenomic Predicted Proteins in Hydroponically and Soil-Grown Tomatoes Rhizosphere.** a) A non-metric multidimensional scaling (NMDS) plot was constructed using Bray-Curtis distances, comparing protein family composition (both annotated and hypothetical) between hydroponic and soil tomato rhizospheres. This visualisation highlights the distinct protein profiles in each cultivation method. b) Volcano plot illustrating significant differences in protein families (log2FoldChange) between hydroponic and soil rhizosphere metagenomes. This plot identifies statistically overrepresented or underrepresented proteins, indicating potential functional adaptations to hydroponic conditions. c) Classification of Clusters of Orthologous Groups (COG) for proteins overrepresented in hydroponic tomato treatments compared to soil.
