## Supplemental Fig. 3 for "Rhizosphere Microbiome Influence on Tomato Growth under Low-Nutrient Settings"

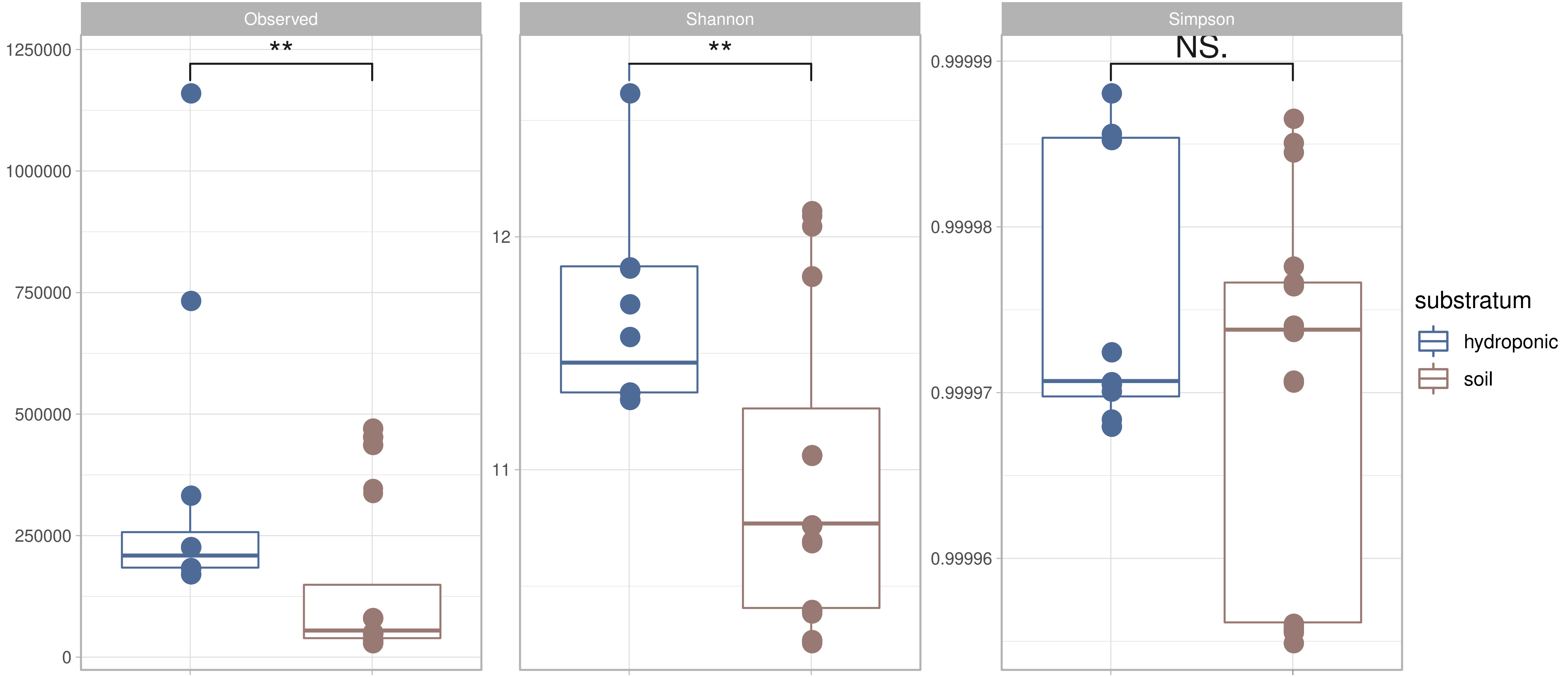


**Figure S3. Metagenomic diversity in hydroponics and soil.** Box plots showing observed protein families, the Shannon and Simpson diversity indexes across hydroponic y soil-grown tomato metagenomes.
