## Supplemental Fig. 4 for "Rhizosphere Microbiome Influence on Tomato Growth under Low-Nutrient Settings"

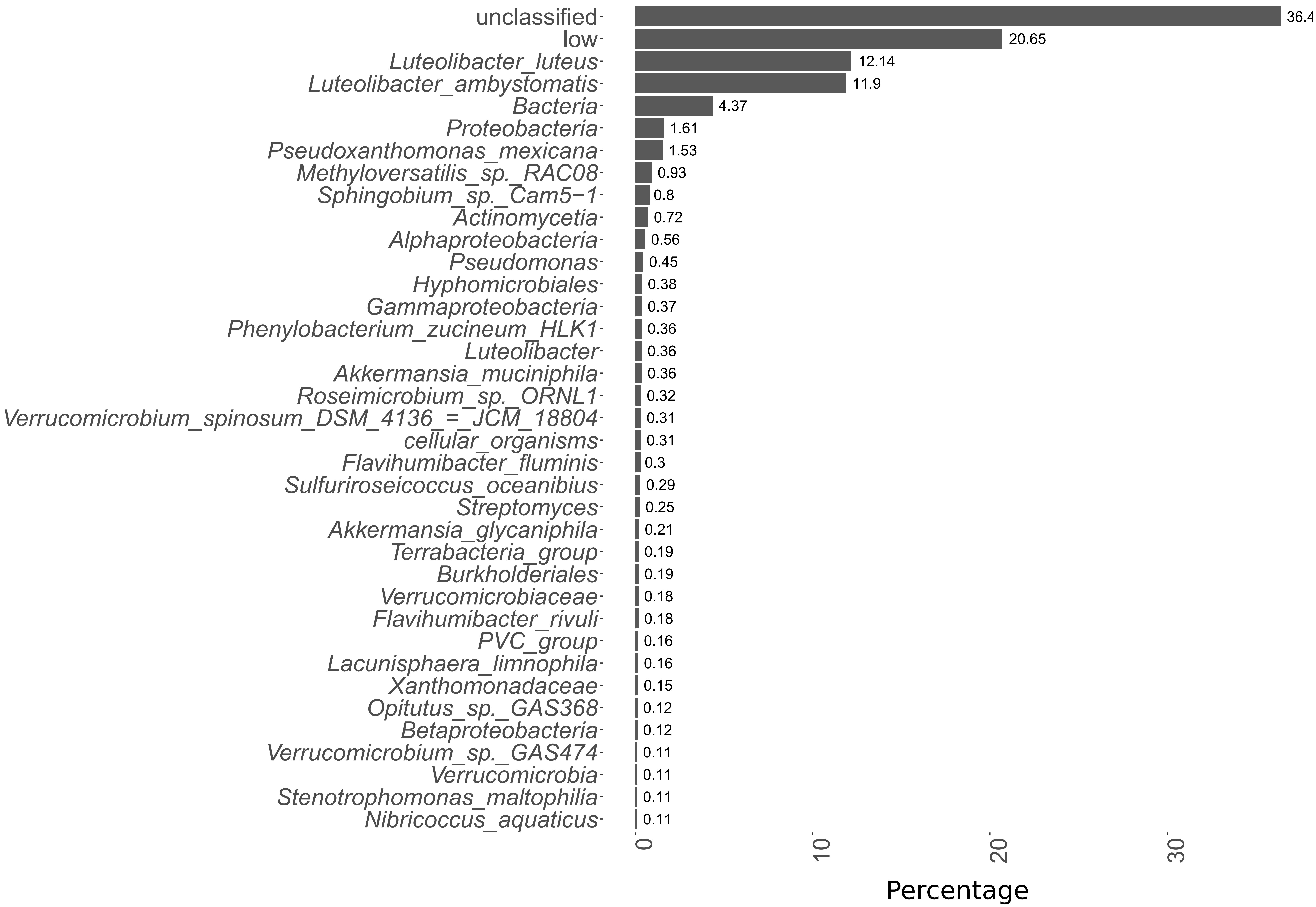


**Figure S4. Taxonomic classification of overrepresented hydroponic proteins.** Percentage of proteins assigned to each taxa (Low percentage ≤ 0.1).
