## Supplemental Fig. 5 for "Rhizosphere Microbiome Influence on Tomato Growth under Low-Nutrient Settings"

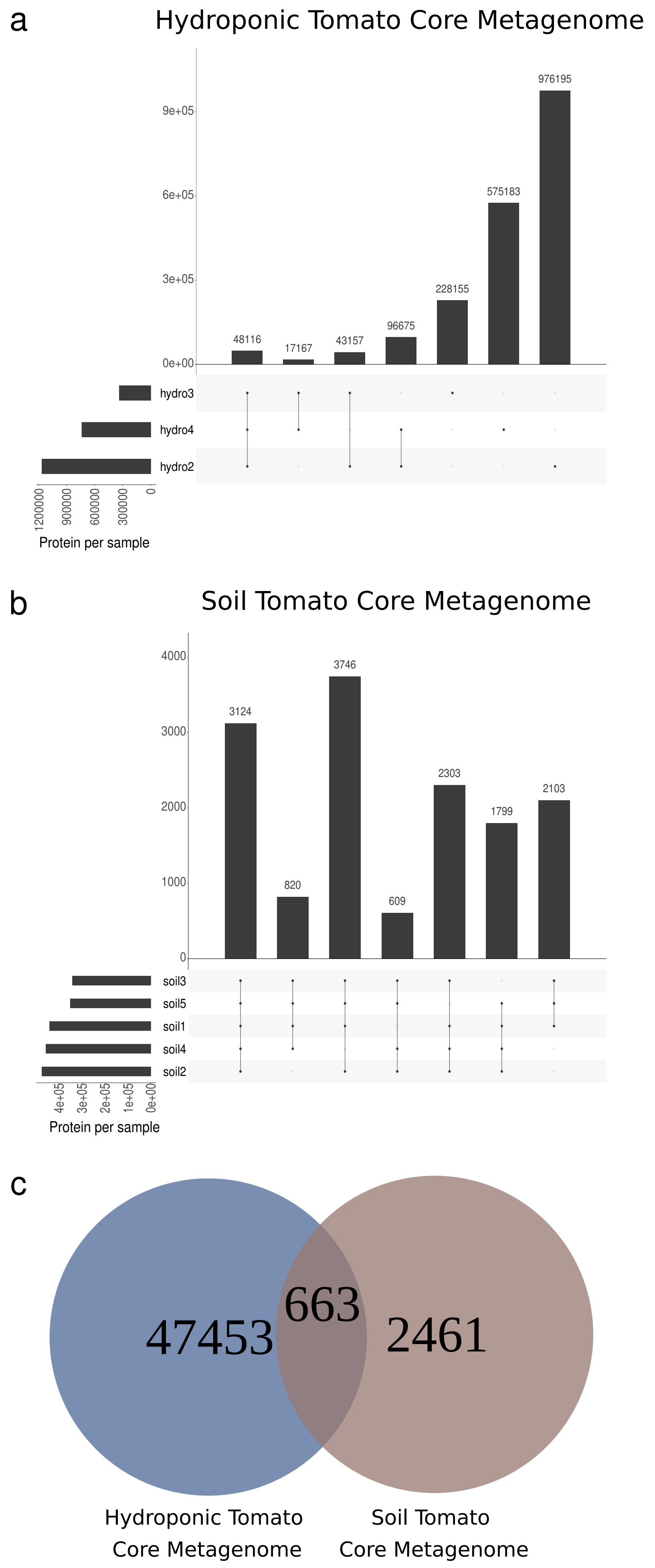


**Figure S5. Tomato core metagenome.** a) Upset plot illustrating the proteins shared by all hydroponic metagenome samples. b) Upset plot showing the proteins found in all soil metagenome samples. c) Venn diagram comparing the hydroponic tomato core metagenome (HTCMe) and the soil-grown tomato core metagenome (STCMe).
